# Contextual Inference Drives Motor Memory Protection and Disruption

**DOI:** 10.64898/2026.08.25.746912

**Authors:** Sumit Sannamath, Rajneet Kaur, Adith Deva Kumar, Adarsh Kumar, Neeraj Kumar

## Abstract

Newly acquired memories are initially fragile and are consolidated into long-term memory over time. Although consolidated memories were once thought to be resistant to interference, accumulating evidence shows that memory reactivation renders them transiently labile, making them susceptible to modification or interference before reconsolidation. Crucially, however, there is substantial variation in whether reactivated memories are disrupted by new information, remain protected from it, or even strengthened by it. The factors that guide this modification are largely unknown. To systematically investigate this, we examined motor memory interference using a classic A-B-A visuomotor rotation paradigm. Participants adapted to a 30-degree clockwise rotation (A) on Day 1. On Day 2, an interfering 30-degree counter-clockwise rotation (B) was introduced under varied conditions: directly without reactivation, after brief reactivation of A, after expression of A without feedback, or following a gradual transition from A to B. The final experiment used explicit contextual cues (a secondary follow-through target) to distinguish A and B trials. Contrary to the simple prediction that reactivation should increase vulnerability to interference, reactivating the original memory before introducing interference protected it, as evidenced by significant savings during relearning on Day 3. In contrast, introducing interference directly, without reactivation, disrupted the original memory. This protection was consistent with a contextual-inference account: the large sensory prediction error experienced during the abrupt transition from A to B served as a latent contextual cue, signaling a new context and thereby shielding the original memory from being overwritten. Eliminating this prediction error through an immediate washout session with a similar error profile or through a gradual A-to-B transition abolished the protective effect and disrupted the original memory. Furthermore, when explicit contextual cues distinguished the two perturbations, memories were protected even in the absence of a salient prediction error. Our findings are consistent with a contextual inference account in which the fate of a consolidated memory, whether it is modified or protected, is shaped by the availability of explicit cues or latent signals such as sensory prediction error at the time when interference is introduced.

## INTRODUCTION

Memory formation is not an instantaneous process. Newly acquired memories are initially fragile and susceptible to disruption. Over time, memories stabilize through a time-dependent stabilization process first characterized by Müller and Pilzecker (1900) called consolidation. During this consolidation window, interference from competing information (Brashers-Krug et al., 1996), electroconvulsive shock (Duncan, 1949), or protein synthesis inhibitors (Agranoff et al., 1966; Agranoff & Klinger, 1964) can impair or erase the memory trace. Once consolidated, these traces were long believed to be stable and resistant to disruption (McGaugh, 2000; Müller & Pilzecker, 1900; Squire & Alvarez, 1995). This classical view, sometimes called the *standard model of consolidation*, held that long-term memory, once formed, is a permanent, fixed engram.

This view was fundamentally challenged by the discovery of memory reconsolidation. Seminal work by Nader and colleagues (2000) demonstrated that reactivating a consolidated fear memory in rodents prior to administration of the protein synthesis inhibitor anisomycin rendered the memory susceptible to erasure similar to post-acquisition amnesia (Nader et al., 2000). This pivotal finding established that retrieval does not merely read out a memory trace, but rather returns that trace to a labile state, requiring *de novo* protein synthesis to stabilize it again. This process is termed as memory reconsolidation (Nader et al., 2000; Nader & Hardt, 2009). Reconsolidation has since been documented across species, brain systems, and memory types, establishing it as a general property of memory rather than a domain-specific anomaly (Dudai, 2006; Lee et al., 2017).

In humans, post-reactivation susceptibility to interference has been demonstrated across multiple memory domains. In procedural motor memory, training on a second finger-tapping sequence immediately after reactivating the first sequence caused retrograde interference with the original sequence, whereas learning without prior reactivation did not (Walker et al., 2003). Analogous effects have been reported for episodic memory, where participants falsely attributed items from an interfering list to the original list, but only when the interfering phase was introduced after explicit reactivation of the original memory (Hupbach et al., 2007). Similarly, distorted recall of action-outcome associations has been observed when interference followed memory reactivation (Sinclair & Barense, 2018). Collectively, these findings suggest that reactivation can open a time-limited window during which consolidated memories in humans are amenable to modification by new experience.

However, reactivation does not invariably render memories vulnerable to disruption. Contrary to prior findings, studies have also found no reliable disruption of a previously learned visuomotor sequence following post-reactivation interference (Hardwicke et al., 2016), failing to replicate the vulnerability effect previously reported (Walker et al., 2003). More strikingly, studies have also reported that interfering experience introduced after reactivation paradoxically strengthened rather than impaired the original motor memory (Herszage & Censor, 2017; Wymbs et al., 2016), a finding that challenges the prediction that reactivation renders memory labile and vulnerable. Further complicating the picture, interference with consolidated motor memories can sometimes occur even in the absence of explicit reactivation (Caithness et al., 2004; Krakauer et al., 2005). The inconsistency across studies in whether reactivation-dependent or reactivation-independent disruption or protection occurs suggests that additional variables govern the fate of a reactivated memory trace beyond the mere occurrence of retrieval.

One compelling candidate mechanism is contextual information. In his influential framework, Bouton (1993) proposed that extinction does not erase the original conditioned association but instead creates a competing memory whose retrieval is context-gated. Contextual cues serve as “occasion setters” that determine which of two co-existing memory representations is expressed in a given situation (Bouton, 1993). Supporting this view, renewal of conditioned responses has been documented in both rodents (Bouton & Bolles, 1979; Bouton & Ricker, 1994) and humans (LaBar & Phelps, 2005; Vansteenwegen et al., 2005) when subjects are returned to the original conditioning environment following extinction. Context-dependent memory formation has also been demonstrated for episodic content. Intrusion errors occurred selectively when interfering learning was introduced in the same spatial context as the original encoding, suggesting that shared context triggers memory updating, whereas context change provides memory protection (Hupbach et al., 2008).

Context need not be explicit for memory protection. Memory protection has been reported even when no overt environmental cues signaled a context change, attributing this protection to implicit sensorimotor variability that participants experienced during the transition between conditions (Wymbs et al., 2016). Such variability generates a mismatch between expected and observed sensory outcomes, called a prediction error, which may function as a latent contextual signal, tagging the new learning episode as arising from a distinct causal state (Exton-McGuinness et al., 2015; Krawczyk et al., 2017). Consistent with this interpretation, prediction error has been shown to facilitate the encoding of non-overlapping memories as distinct traces (Greve et al., 2017; Henson & Gagnepain, 2010). The COntextual INference (COIN) model (Heald et al., 2021) formalizes this intuition. The brain continuously performs Bayesian inference over latent contexts, updating an existing memory only when the inferred context matches that of prior learning, and creating a new memory trace when it does not. The prediction error, therefore, could serve as the critical signal driving context differentiation. Importantly, prediction error has also been shown to destabilize existing memories by triggering reconsolidation-mediated updating (Sevenster et al., 2013; Sinclair & Barense, 2018, 2019), suggesting that its role in memory modification may depend critically on when during the memory episode it occurs, at retrieval, or at the onset of new interfering learning.

Despite considerable progress, there is no concrete picture of the driver of memory protection or modification. The conditions that determine whether interference following reactivation disrupts, protects, or strengthens a motor memory remain poorly understood. The existing literature is characterized by substantial cross-study variability in paradigm design, including the presence or absence of explicit context cues, the nature and timing of interfering perturbations, and the use of reactivation versus expression probes, making direct comparisons difficult. The present study addresses this gap by systematically examining these factors within a unified visuomotor rotation framework using the classical A-B-A interference design. Across experiments, we assessed how the modification or protection of an original motor memory by an interfering visuomotor perturbation is influenced by (i) whether the memory is reactivated, not reactivated, or merely expressed prior to interference, and (ii) whether explicit or latent contextual cues are available to differentiate the learning episodes. Our results demonstrate that both explicit and latent contextual information shape reconsolidation outcomes. In the absence of disambiguating contextual signals, interference preferentially disrupts the original memory, whereas the presence of such signals, whether overt or implicit, can lead to motor memory protection.

## METHODS

### Participants

A total of 127 right-handed, healthy individuals (86 men, 41 women; age = 22.67 ± 3.45; age range = 18-33 years) participated in the study. The Edinburgh Handedness Inventory was used to assess their handedness (Oldfield, 1971). Participants reported no history of neurological, psychological, or any physical problems that could affect their performance of the task. All participants provided written informed consent and were compensated for their time. All the experimental procedures were approved by the institute’s ethics committee.

### Power Analysis

A priori power analysis was conducted using G*Power 3.1 (Faul et al., 2007) to estimate the required sample size for detecting differences between two dependent means (matched pairs). Assuming an effect size of (d_z_ = 0.9), an alpha level of 0.05, and a statistical power of 0.80, the analysis indicated that a minimum total sample size of 10 participants would be required to detect the expected effect. This sample size is consistent with previous studies employing the visuomotor adaptation paradigm (Kitago et al., 2013; Morehead et al., 2015; Taylor & Ivry, 2011).

### Study Apparatus

The experimental setup comprised a virtual reality system in which the participant sat in front of a digitizing tablet (GTCO CalComp) and used a stylus to make planar movements on the tablet. The hand’s position was displayed as a cursor on a monitor mounted above the tablet. The view of the hand was blocked by placing a mirror between the screen and the tablet, which reflected the screen, showing the hand’s movement as a cursor on the mirror (Figure 1A). The position of the cursor can be veridical or distorted relative to the motion of the hand. Hand movement data was recorded at 120 Hz.

**Figure 1:**
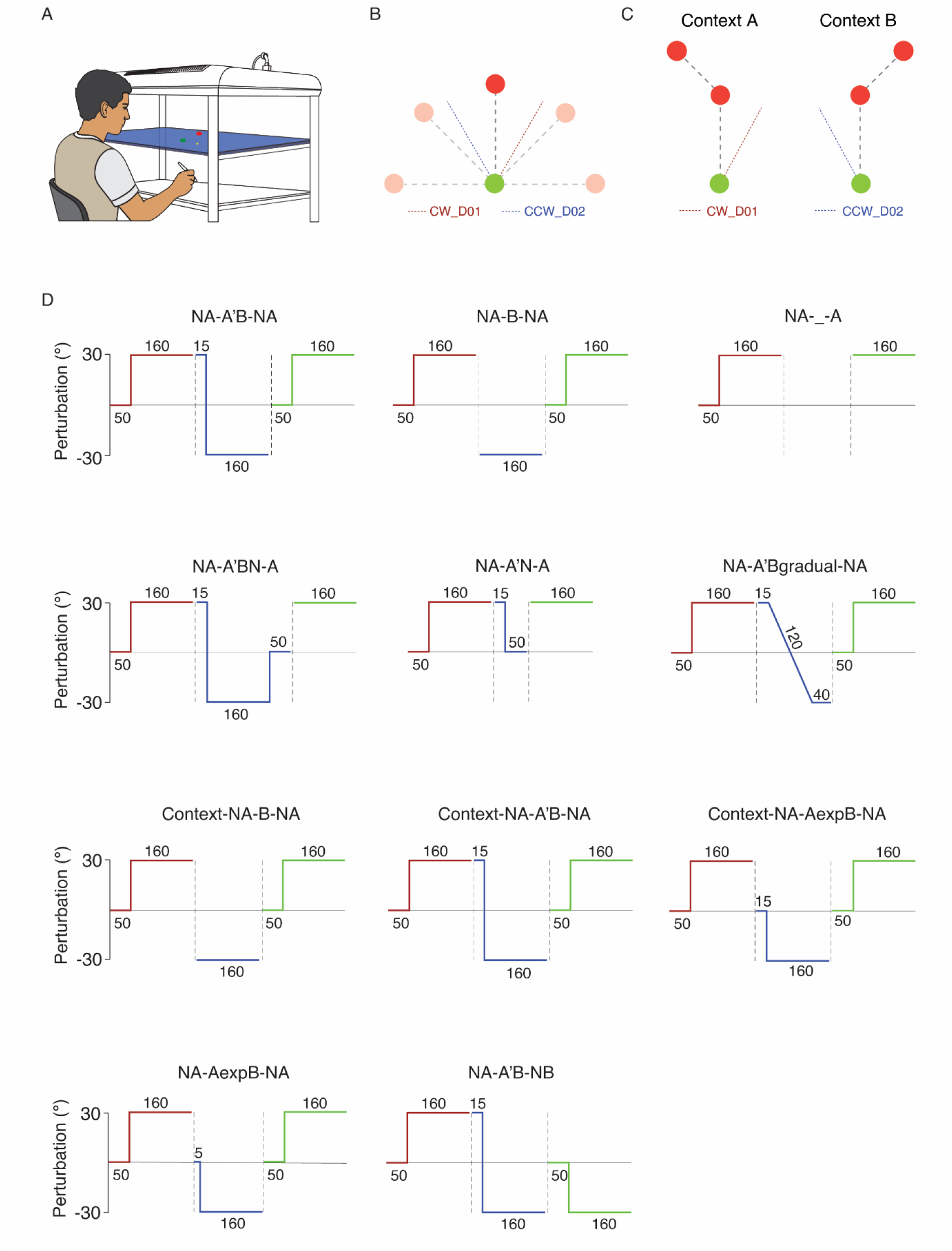
Illustration of the experimental setup and experiment design **(A)** A virtual reality system, where participants viewed the start circle (green) and target circle (red) projected from the monitor on a semi-silvered mirror placed between the monitor and digitizing tablet on which the participants made reaching movements from the start circle to the target circle using a stylus. Participants received veridical or perturbed feedback about hand position through a cursor (yellow dot) displayed on the screen. **(B)** Schematic diagram showing the five target positions (0°, 45°, 90°, 135°, and 180°) used in the no explicit context groups. **(C)** Schematic representation of the target (90°) and contextual cue (secondary follow-through target at 135°-A context and 45°-B Context) used in the explicit context groups. The applied cursor rotations are represented using colored dotted lines, with the red dotted line indicating the 30° clockwise rotation introduced on Day 1 (CW_D01), and the blue dotted line indicating the interfering 30° counter-clockwise rotation introduced on Day 2 (CCW_D02). **(D)** Trial structure across different groups: on Day 1 (red), all subjects, regardless of group, began with 50 no-rotation baseline trials (N), followed by 160 trials of adaptation to a 30° clockwise (CW) rotation (A). **First row:** Group NA-A’B-NA (left panel) - Day 2 (blue) began with brief re-exposure to the 30° CW rotation for 15 trials (A’), immediately followed by 160 trials of 30° CCW rotation (B). Similar to Day 1, Day 3 (green) began with 50 no-rotation trials (N), to wash out any effects of anterograde interference, followed by 160 trials of 30° CW rotation (A). Group NA-B-NA (middle panel) - Day 2 (blue) consisted of 160 B-trials of 30° CCW rotation, with no prior re-exposure to CW rotation. Day 3 was the same as Day 1. Group NA-_-A (right panel) - there was a 48-hour break after Day 1, then on Day 3, the 30° CW rotation was reintroduced for 160 A-trials. **Second row:** Group NA-A’BN-A (left panel) - Day 2 began with 15 trials of re-exposure to 30° CW rotation(A’), immediately followed by 160 trials of 30° CCW rotation (B), further followed by 50 no-rotation trials (N). Day 3 consisted of 160 trials of 30° CW rotation (A). Group NA-A’N-A (middle panel) - Day 2 began with 15 trials of 30° CW(A’), immediately followed by 50 no-rotation trials (N) with no exposure to the interfering CCW rotation. Day 3 consisted of 160 trials of 30° CW rotation (A). Group NA-A’Bgradual-NA (right panel) - Day 2 began with 15 trials of 30° CW rotation (A’), followed by a gradual introduction of 30° CCW rotation over 120 trials, then 40 trials at full 30° CCW (Bgradual). After a 24-hour gap, Day 3 began with 50 no-rotation trials (N) followed by 160 trials of 30° CW rotation (A). **Third row:** The trial structure for the first two context groups - Context-NA-B-NA (left panel) and Context-NA-A’B-NA (middle panel) was same as the corresponding first two groups (First row), with the only difference being for group Context-NA-AexpB-NA (right panel) - Day 2 began with 15 error clamped A context trials (Aexp), that is, the cursor path was clamped to 90° irrespective of hand movement, immediately followed by 30° CCW rotation for 160 trials (B). Day 3 (green) began with 50 no-rotation trials (N) to wash out any effects of anterograde interference, followed by 160 trials of 30° CW rotation (A). **Fourth row:** Group NA-AexpB-NA (left panel) - Day 2 began with 5 no cursor feedback trials (Aexp), immediately followed by 30° CCW rotation for 160 trials (B). After a gap of 24 hours, Day 3 began with 50 no-rotation trials (N) followed by 160 trials of 30° CW rotation (A). Group NA-A’B-NB (middle panel) began Day 2 with fifteen 30° CW rotation trials (A), immediately followed by 160 trials of 30° CCW rotation (B). After a gap of 24 hours, Day 3 began with 50 no rotation trials (N), which was immediately followed by 160 trials of 30° CCW rotation (B), which they learned on Day 2.

### Task Procedure

#### Motor Learning Task

The participants performed 14-cm-long reaching movements from a central start circle (green circle, diameter 8 mm) to five radially arranged targets at 0°, 45°, 90°, 135°, and 180° (red circle, diameter 8 mm) (Figure 1B). After familiarization with the experimental setup, participants were instructed to move the yellow cursor (dot, diameter 3 mm) to the starting point and wait for 250 ms for the auditory “beep” cue. Upon presentation of the auditory cue, a target appeared at one of five possible locations.

Participants were instructed to make a ballistic movement towards the presented target, reach within 2 s, and maintain their position at the target until an auditory cue signaled the end of the trial. At the end of the trial, they received numerical feedback based on their performance on the trial. The feedback screen was presented for 500 ms on the screen, ranging from 10 to 0 points based on the cursor’s distance from the target, where 10 indicates the cursor was inside the target. Points did not influence the compensation provided to the participants and were not analyzed either. The order of presentation of the target was predetermined pseudo-randomly before the experiment, so that the N^th^ trial would be the same for all participants.

Baseline sessions (N) were performed by each participant, during which no perturbations were applied to the cursor feedback to establish participants’ initial baseline performance under accurate visual feedback. Learning sessions consisted of either a 30° clockwise (CW) perturbation (A-trials) or a 30° counterclockwise (CCW) perturbation (B-trials). Throughout the experiment, catch trials were intermittently introduced during which the cursor was not displayed, while participants were instructed to perform the movement as they had during the previous trial. These catch trials were used to assess the expression of the learned visuomotor perturbation without visual feedback.

#### Motor Learning Task with explicit Contextual Cues

The participants made reaching movements from the starting circle (blue, diameter 8 mm) towards the primary and secondary target circles (red, diameter 8 mm). The primary target was located 12 cm straight to the start point, while the secondary follow-through target was positioned 45° to the left (context cue A for A (CW) perturbation) or to the right (context cue B for B (CCW) perturbation), 8 cm away from the primary target (Figure 1C).

After the familiarization and practice session, participants were asked to bring the yellow cursor dot (3 mm diameter) to the starting blue circle and wait for 500 milliseconds for the circle to change its color to green. As the starting circle changes color to green, the central primary target and the secondary target (either on the left or the right) will appear simultaneously on the screen (Howard et al., 2013; Kumar et al., 2026; Sheahan et al., 2016). The participants were instructed to make a swift single movement to the secondary target, ensuring they pass through the primary target, and to hold their position on the secondary target until they hear an auditory “beep” cue indicating the end of the trial. At the end of each trial, participants received a feedback screen displaying a numerical performance score along with feedback on their movement velocity. The numerical score ranged from 0 to 10 points and was determined by the distance of the cursor from the target, with a score of 10 indicating that the cursor was positioned within the target. Participants also received feedback on their movement velocity, displayed as “Too Slow,” “Good,” or “Too Fast.” The feedback screen was presented for 500 ms before the next trial began. They were instructed to maintain a “Good” movement velocity throughout the experiment. If the participants failed to pass through the central red circle, they received “Incorrect Trial” feedback and did not receive any points for that trial. The points did not influence the participants’ compensation and were not included in the data analysis.

During the baseline session, pseudo-randomized A and B context trials were presented without any perturbation to establish participants’ initial baseline performance under accurate visual feedback. During the learning sessions, each trial was associated with a specific perturbation based on the contextual cue. If the secondary target was presented on the left side (cue A), the perturbation was 30° clockwise, and if it was presented on the right side (cue B), the perturbation was 30° counterclockwise. Thus, the secondary target served as an explicit contextual cue indicating the perturbation associated with each trial. Throughout the experiment, catch trials (no visual feedback) were interspersed to assess the expression of learning in the absence of visual error feedback. Participants were not provided with any explicit description regarding the nature of the secondary targets and their relation to the perturbations.

### Experimental Design

#### Experiment-1

The experiment was designed to explore the effect of interfering learning on consolidated motor memory. We hypothesized that reactivation of a previously consolidated memory prior to exposure to an interfering perturbation would render the memory susceptible to disruption.

To test this, participants were randomly assigned to one of the three experimental groups that differed in what occurred on Day 2 and Day 3, while the protocol on Day 1 remained the same across groups. All the participants first completed a 50-trial veridical-feedback session (N) to establish baseline reaching performance, followed by a 160-trial learning session in which they adapted to a 30° CW visuomotor rotation (A) on Day 1.

On Day 2, group *NA-A’B-NA* was briefly re-exposed to a 30° CW rotation (A’) over 15-trials to reactivate the previously consolidated A memory. This was immediately followed by a 160-trial interference session with the opposing 30° CCW rotation (B). On Day 3, the participants were provided with a veridical-feedback washout session (50 N-trials) to eliminate any anterograde carry-over of B from Day 2. This was followed by a 160-trial relearning session with the A rotation. This group was designed to test whether reactivation prior to interference renders the A memory vulnerable to disruption (Figure 1D, first row, left panel).

Group *NA-B-NA* received the same 160-trial B interference session on Day 2 without any preceding A trials, allowing us to determine whether retrograde interference from B-learning can disrupt a consolidated memory even in the absence of prior reactivation of the initial memory. The Day 3 protocol was identical to that of the *NA-A’B-NA* group (Figure 1D, first row, middle panel).

Group *NA-_-A* served as a no-interference control, receiving no Day 2 exposure and proceeding directly to the Day 3 relearning session (160 A-trials). This group provided a pure estimate of savings in the absence of interfering learning, against which the other two groups’ savings could be compared (Figure 1D, first row, right panel).

#### Experiment-2

Experiment 1 established that reactivation prior to interference protects the consolidated A memory, which we attributed to the abrupt sensory prediction error at the A-to-B transition serving as a latent context-change signal. However, two explanations remain possible. Reactivation may reflect a general resistance to overwriting, such that any interfering information introduced after reactivation fails to disrupt the original memory regardless of how it is introduced. Alternatively, protection may depend specifically on the prediction error generated at the transition into the interfering rotation, rather than on reactivation itself. To dissociate these possibilities, we asked what happens to A memory when the interfering B learning is returned to baseline immediately after acquisition, before it has any opportunity to consolidate. Under the general-resistance account, eliminating B before consolidation should produce equivalent or stronger protection of A, since the interfering memory never had the opportunity to compete with the original. Under the prediction-error account, whether B consolidates should be irrelevant; what matters is the prediction error available to the motor system at each perturbation transition, including the washout.

To test this, we introduced the *NA-A’BN-A* group, in which participants reactivated the consolidated A memory (15 A-trials) and were then exposed to the interfering B rotation (160 B-trials), which was immediately followed by a washout session (50 N-trials) on Day 2, thereby preventing its consolidation (Figure 1D, second row, left panel).

We additionally introduced a control group, *NA-A’N-A*, in which no interfering rotation was presented at all, and the reactivated A memory was followed directly by a washout session, allowing us to isolate the effect of no-perturbation trials on a reactivated memory in the complete absence of interference. The participants in group *NA-A’N-A* performed the reactivation session (15 A-trials) followed by the washout session (50 N-trials) without any interference on Day 2 (Figure 1D, second row, middle panel). On Day 1, both groups performed a baseline session (50 N-trials) followed by the learning session (160 A-trials). On Day 3, both groups performed only the relearning session (160 A-trials).

#### Experiment-3

Experiments 1 and 2 suggest that the sensory prediction error experienced at perturbation transitions is a plausible driving signal for motor memory protection. Experiment 3 was designed to test this directly by eliminating the abrupt prediction error at the A’-to-B transition on Day 2 altogether. If prediction error at the transition serves as a critical contextual signal driving separate memory encoding, then removing it, by introducing the interfering rotation gradually rather than abruptly, should abolish memory protection even when reactivation precedes interference. To test this, we introduced group *NA-A’Bgradual-NA* to remove the large prediction error experienced during the transition from A’ to B perturbation on Day 2. Similar to experiments 1 and 2, participants performed a baseline session (50 N-trials) followed by the learning session (160 A-trials) on Day 1. On Day 2, participants performed a reactivation session (15 A-trials), followed by a gradual transition from 30° CW (A) towards 30° CCW (B) at an increment of 0.5° across 120 trials. The perturbation was held constant at 30° CCW for an additional 40 trials. On Day 3, participants performed a washout session (50 N-trials) to eliminate any anterograde effects of learning the B perturbation. This was followed by a relearning session (160 A-trials) (Figure 1D, second row, right panel).

A critical corollary of the prediction-error account is that when an abrupt transition error is present, and context differentiation succeeds, as in Experiment 1, the interfering B rotation could be encoded as a distinct memory and remains independently retrievable. To examine this possibility, we introduced the *NA-A’B-NB* group, which underwent the identical Day 2 protocol as *NA-A’B-NA* in Experiment 1, reactivation of A followed by 160-trial B-learning but was retested on B rather than A on Day 3 (Figure 1D, fourth row, middle panel).

#### Experiment-4

This experiment was designed to determine whether an explicit contextual cue, rather than an experienced prediction error, can drive the distinct encoding of A and B memories. Specifically, we asked whether tagging each perturbation with a distinct contextual cue would protect the original A memory from interference, and whether such protection would extend even to the condition that produced retrograde interference in the *NA-B-NA* group in Experiment 1. To address this, we replicated the two core groups from Experiment 1, *NA-B-NA* and *NA-A’B-NA*, but tagged each rotation with a distinct explicit contextual cue in the form of a secondary follow-through target, yielding groups *Context-NA-B-NA* and *Context-NA-A’B-NA*, respectively.

Similar to Experiment 1, participants performed a baseline session (50 N-trials) followed by the learning session (160 A-context trials with A perturbation) on Day 1. The participants in *Context-NA-B-NA* group directly performed the interference session (160 B-context trials with B perturbation) on Day 2 (Figure 1D, third row, left panel), while the participants in *Context-NA-A’B-NA* group performed the reactivation session (15 A-context trials with A perturbation) before the interference session (160 B-context trials with B perturbation) (Figure 1D, third row, middle panel). On Day 3, participants in both groups performed a washout session before the relearning session (160 A-context trials with A perturbation).

#### Experiment-5

For the protection observed in the initial three experiments, the contextual signal arose from error-driven retrieval of A memory, immediately preceding the interfering B rotation. Experiment 5 was designed to determine whether error-driven retrieval is necessary for contextual change, or whether passive expression of the consolidated memory in the complete absence of error feedback is sufficient.

To test this, we introduced two groups: *NA-AexpB-NA* (Figure 1D, fourth row, left panel) and *Context-NA-AexpB-NA* (Figure 1D, third row, right panel). As in previous experiments, Day 1 for both groups began with a 50-trial veridical-feedback baseline session, followed by a 160-trial learning session to adapt to a 30° CW rotation (A). On Day 2, participants performed a brief expression session consisting of 5 no-cursor feedback trials in the *NA-AexpB-NA* group, and 15 A-context trials with cursor feedback clamped to zero (error-clamped trials) in the *Context-NA-AexpB-NA* group. Following the respective expression session, both groups were exposed to interfering rotation (160 B-trials). On Day 3, participants in both groups performed a washout session (50 N-trials) to remove any anterograde effects of B learning followed by a 160-trial relearning session (A).

### Data Analysis

The data was collected and pre-processed using MATLAB (version 9.13.0, R2022b; The MathWorks Inc., 2022). Hand position data from each trial were recorded for offline analysis. Kinematic data were filtered using a low-pass Butterworth filter with a cutoff frequency of 8 Hz, and instantaneous velocity was derived from the filtered position data. The primary dependent measure was direction error (DE), defined as the angular deviation between the cursor position at peak tangential velocity and the straight-line path from the start position to the target. Direction errors were baseline-corrected by subtracting each participant’s mean direction error across the first 50 baseline trials (N) on Day 1. Catch trials were excluded from all analyses. Trials were grouped into bins of five consecutive trials. Learning within each session was characterized by comparing the early and late phases, as indicated by the DE of the first three and last three bins, respectively. For the reactivation session, DE on its last bin was compared with DE on the last bin of learning to indicate successful reactivation of A memory. For the expression session, the no-cursor feedback bin in the no-context group was compared with zero, whereas in the context group, the first and last bins of the expression session were compared to confirm successful reactivation. Savings on Day 3 were defined as a significantly lower direction error during the early phase of the relearning session on Day 3 relative to the early phase of the learning session on Day 1, reflecting faster reacquisition of the perturbation.

Across all experiments, five participants were excluded from analysis for failing to complete all required sessions. For the remaining 122 participants, individual trials were flagged as bad trials and excluded if the participant failed to initiate movement, did not complete the movement within the required time window, or lifted the pen from the tablet mid-trial, resulting in data loss. Across all included participants, bad trials accounted for 0.38% of all trials. All statistical tests are reported alongside the results to which they pertain. All statistical analyses were performed using RStudio (version 2025.9.2.418; Posit team, 2025). The significance threshold was set at α = 0.05 throughout, and all the post hoc comparisons were corrected using the Benjamini-Hochberg (BH) method. Bayesian analyses were additionally performed to corroborate each null result, with Bayes factors (*BF*₁₀) computed using the BayesFactor package in R.

## RESULTS

### Experiment 1: Reactivation of consolidated memory protects it against subsequent interference

To investigate how opposing interfering information affects a previously consolidated motor memory, and whether the fate of that memory depends on its reactivation prior to interference, we assigned participants to three groups: *NA-A’B-NA*, NA-B-NA, and *NA-_-A*. On Day 1, all groups first performed reaching movements under veridical cursor feedback to establish a baseline performance before adapting to the 30° CW visuomotor rotation (A). Participants across groups showed a typical canonical reduction of direction error across early and late learning session (Figure 2). A mixed ANOVA showed significant reduction in the direction error from early (Mean ± SD = 25.02 ± 4.76°) to late (Mean ± SD = 6.50 ± 4.17°) session [*F*(1, 33) = 865.54, *p* < 0.001, ω² = 0.81] and confirmed equivalent acquisition of A with no significant between-group difference [*F*(2, 33) = 1.04, *p* = 0.366, ω² = < 0.01] with support from the Bayesian analysis against group effect (*BF*_10_ = 0.51).

**Figure 2:**
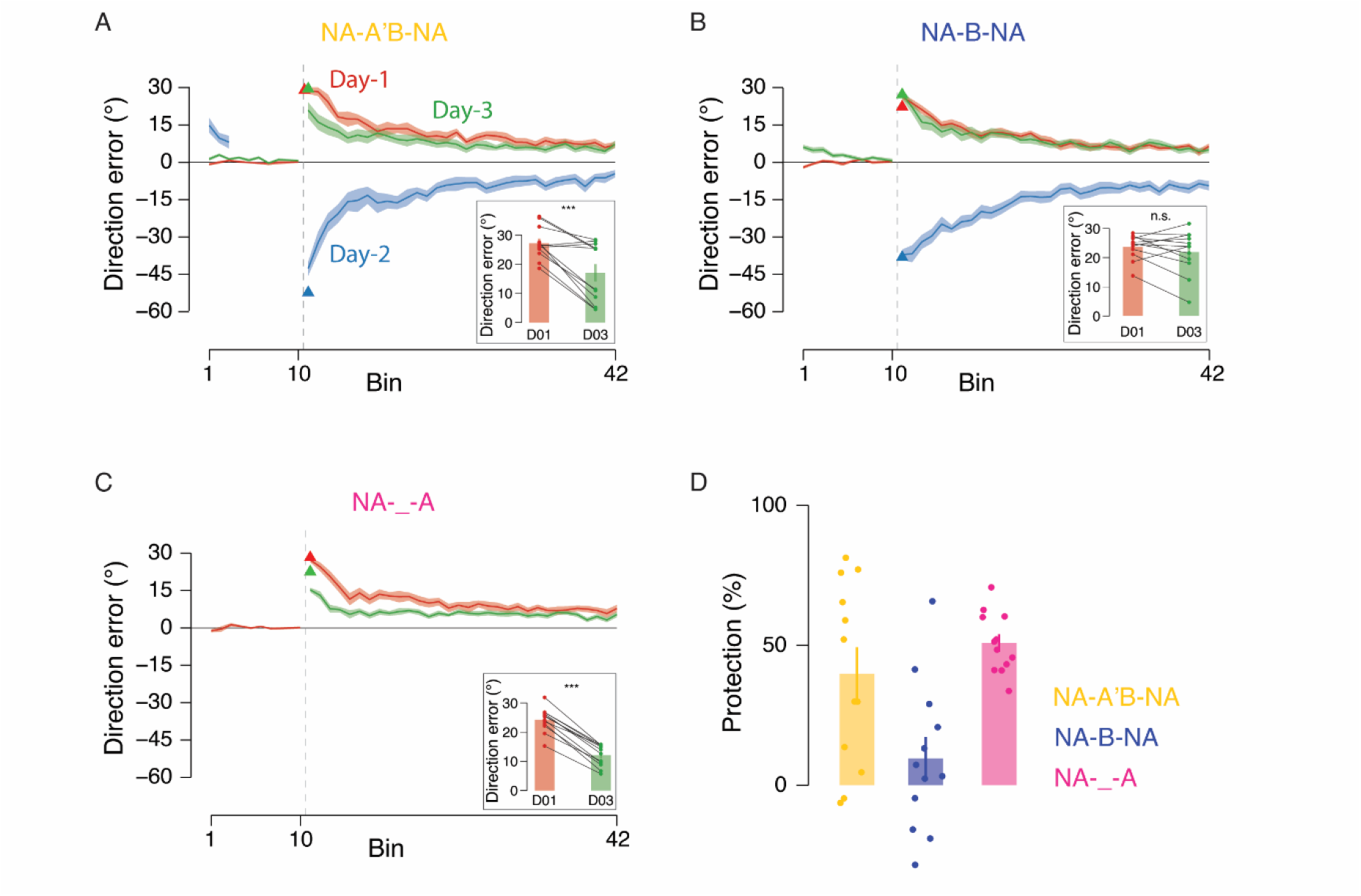
Reactivating the consolidated motor memory prior to interference protected it against disruption. Change in mean direction error (°) across bins for participants in **(A)** group NA-A’B-NA, **(B)** group NA-B-NA, and **(C)** group NA-_-A. Shaded ribbons indicate ±SE across participants. Triangles mark the first trial of the Day 1 learning, Day 2 interference, and Day 3 relearning sessions. Red, blue, and green colors denote Day 1, 2, and 3, respectively. All three groups showed equivalent acquisition of the 30° CW rotation (A) on Day 1. On Day 2, both NA-A’B-NA and NA-B-NA groups fully adapted to the opposing 30° CCW rotation (B) to equivalent asymptotic levels. On Day 3, the NA-A’B-NA group, which reactivated A memory immediately before B-learning, relearned A significantly faster than on Day 1, demonstrating robust savings and confirming that the consolidated A memory was protected despite full adaptation to the opposing rotation. The NA-_-A group, which received no interference on Day 2, similarly showed strong savings, establishing the upper bound of memory retrieval in the absence of any competing experience. By contrast, the NA-B-NA group, which received identical B-learning on Day 2 but without prior reactivation of A, showed no savings and relearned A as naive on Day 1, indicating complete disruption of consolidated memory. Insets: Average direction error (°) across the first three bins on Day 1 (red) and Day 3 (green) for each participant (dots), connected by lines to illustrate individual-level change. **(D)** Protection index [(Day 1 early error − Day 3 early error) / Day 1 early error × 100%] for each group, quantifying the degree to which the A memory was retained across the three-day interval. A value of 100% indicates complete retention; values near zero indicate disruption of the A memory. The NA-B-NA group had a significantly lower protection index than both NA-A’B-NA and NA-_-A, while the latter two groups did not differ from one another, confirming that disruption in NA-B-NA reflects retrograde competition from B-learning rather than the passage of time. Individual participant values are shown as dots.

On Day 2, participants in the *NA-A’B-NA* group underwent a brief reactivation session (15 A-trials) of the original CW rotation. Direction error at the end of this session did not differ significantly from the last bin of Day 1 learning [*t*(11) = 0.61, *p* = 0.552, Cohen’s *d*_z_ = 0.19 ] with strong support for the null hypothesis from the Bayesian analysis (*BF*_10_ = 0.34), confirming that the previously consolidated A memory was successfully accessed and restored to its prior performance level. Both *NA-A’B-NA* following reactivation and *NA-B-NA* fully adapted to the opposing 30° CCW rotation across the interference session [*F*(1, 22) = 220.57, *p* < 0.001, ω² = 0.75], reaching equivalent asymptotic performance with no significant between-group difference at end of B training [*F*(1, 22) = 1.37, *p* = 0.255, ω² < 0.01] with support from the Bayesian analysis against group effect (*BF*_10_ = 0.61).

On Day 3, both groups began by performing the reaching movement under veridical feedback to wash out any anterograde effects of learning the B perturbation. The *NA-_-A* group, which had received no interfering perturbation on Day 2, did not undergo a washout. We examined, within each group, whether participants exhibited savings, reflected in faster relearning relative to their Day 1 acquisition. The participants in the *NA-A’B-NA* group relearned the A perturbation faster on Day 3 in comparison to their initial learning on Day 1 and showed significant savings [*t*(11) = 4.54, *p* < 0.001, Cohen’s *d*_z_ = 1.37] (Figure 2A). Similar savings were observed in the *NA-_-A* group, where there was no interference on Day 2 [*t*(11) = 15.64, *p* < 0.001, Cohen’s *d*_z_ = 4.71] (Figure 2C). However, the *NA-B-NA* group showed no reliable savings [*t*(11) = 1.23, *p* = 0.245, Cohen’s *d*_z_ = 0.37] with support for the null hypothesis from the Bayesian analysis (*BF*_10_ = 0.53) (Figure 2B). Participants relearned the A perturbation on Day 3 as naïve and similar to their Day 1 learning.

To quantify the degree of memory protection across groups, we computed a protection index for each participant, defined as the percentage reduction in early-phase direction error from Day 1 to Day 3 A-learning, relative to Day 1 error [(Day 1 early error − Day 3 early error) / Day 1 early error × 100%]. A value of 100% indicates complete retention; values close to zero indicate little or no protection of the A memory. A one-way ANOVA on the protection index revealed a significant group effect [*F*(2, 33) = 8.61, *p* < 0.001, ω² = 0.30] (Figure 2D). Post-hoc comparisons confirmed that the protection index was significantly lower in the *NA-B-NA* group than in *NA-A’B-NA* (*p* = 0.009), and *NA-_-A* (*p* = 0.001). No significant differences were observed between groups where either interference was introduced post-reactivation on Day 2 or when there was no interference at all (*p* = 0.291). These results confirm that the interference-induced disruption observed in the *NA-B-NA* group was not driven by passage of time, which would have affected the *NA-_-A* group equally. The selective disruption in *NA-B-NA* therefore reflects retrograde competition specifically arising from B-learning.

Collectively, the results of Experiment 1 demonstrate that the fate of a consolidated motor memory following interference is not determined by the mere presence of competing information, but is critically modulated by whether the original memory was accessed immediately prior to that interference. Prior access through error-based reactivation was associated with robust retention of the A memory despite full adaptation to the opposing B rotation. Notably, this outcome is contrary to our initial hypothesis and to the classical reconsolidation prediction that reactivation should increase vulnerability to interference. Rather than being disrupted, the reactivated memory was protected. One plausible reason for this dissociation is that accessing the A memory prior to B-learning generates a sensory prediction error at the A’ to B transition, which functions as a latent contextual signal. This signal may drive the motor system to infer that B arises from a distinct causal context, triggering separate encoding of B perturbation rather than updating the memory of A (Exton-McGuinness et al., 2015; Heald et al., 2021; Krawczyk et al., 2017). In the *NA-B-NA* group, the absence of any prior A-access means no such prediction error is generated at the onset of B; the two opposing rotations therefore compete within a shared representational context, and B successfully overrides the original A memory. This was also supported by the observation that participants learned the B perturbation faster in the *NA-A’B-NA* than in the *NA-B-NA* group. Participants in the *NA-A’B-NA* group took approximately 7 bins to reach 67% of the end of B learning compared to 11 bins in the *NA-B-NA* group. This faster learning was achieved despite the larger error in the first trial, which was due to the reactivation session. This would be expected if participants were considering B perturbation as a new context and learned it with reduced anterograde interference.

### Experiment 2: Washout does not necessarily erase the memory preceding it

Experiment 2 was designed to test whether washing out the interfering B rotation before it could consolidate would further enhance the protection of initial A memory, beyond what was observed in the *NA-A’B-NA* group of Experiment 1. To address this, we introduced the group *NA-A’BN-A*. As in Experiment 1, participants began Day 1 with a veridical-feedback baseline session, followed by acquisition of the 30° CW rotation (A). Participants learned the A perturbation with a significant reduction of direction error from the early (Mean ± SD = 21.11 ± 5.79°) to the late phase (Mean ± SD = 6.23 ± 2.26°) of the learning session [*t*(11) = 8.49*, p* < 0.001, Cohen’s *d_z_* = 2.56].

On Day 2, participants first reactivated the consolidated A memory under 30° CW perturbation. Direction error at the end of the reactivation session did not differ from the last bin of Day 1 learning [*t*(11) = 1.46, *p* = 0.172, Cohen’s *d*_z_ = 0.44]. Bayesian analysis further supported the null hypothesis (*BF*_10_ = 0.67), confirming successful reactivation. Post reactivation, participants successfully adapted to the interfering 30° CCW perturbation [*t*(11) = 9.86, *p* < 0.001, Cohen’s *d*_z_ = 2.97]. This was immediately followed by no-perturbation trials (N), which returned performance to baseline [*t*(11) = 6.73, *p* < 0.001, Cohen’s *d*_z_ = 2.03]. Since the interfering learning was washed out immediately without any opportunity for consolidation, we expected strong savings of A learning on Day 3. Contrary to expectations, we found no savings for A learning on Day 3 [*t*(11) = 1.80, *p* = 0.099, Cohen’s *d*_z_ = 0.54]. Bayesian analysis provided further support for this null effect (*BF*_10_ = 0.99). Instead of washing out learning of B perturbation, the N-trials seemingly washed out or modified the A memory (Figure 3A).

**Figure 3:**
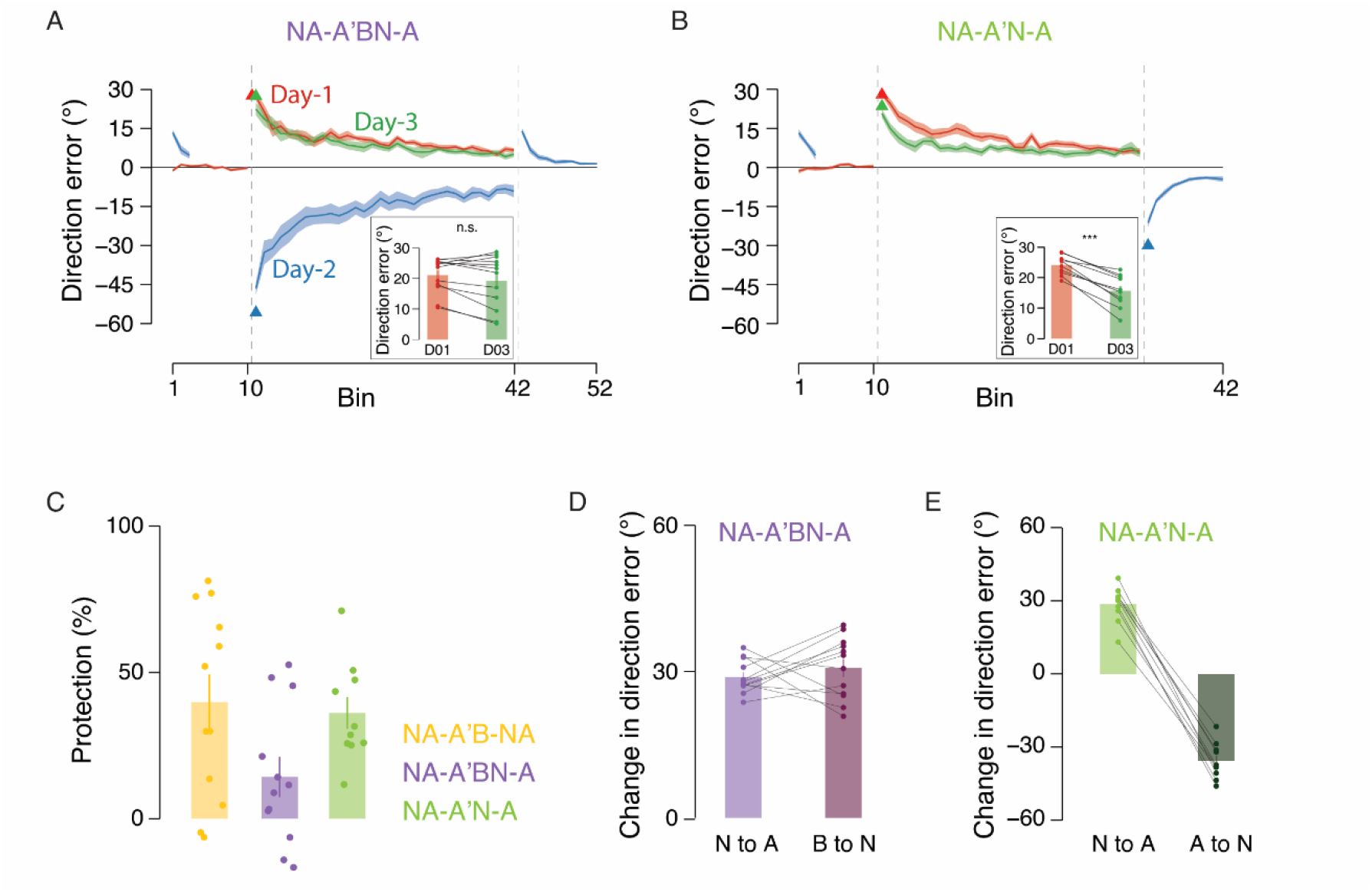
The direction of the sensory prediction error at perturbation transitions determined the fate of the consolidated motor memory. Change in mean direction error (°) across bins for participants in **(A)** group NA-A’BN-A and **(B)** group NA-A’N-A. Shaded ribbons indicate ±SE across participants. Triangles mark the first trial of the Day 1 learning, Day 2 interference or washout, and Day 3 relearning sessions. Red, blue, and green colors denote Day 1, 2, and 3 respectively. Both groups showed successful acquisition of the 30° CW rotation (A) on Day 1 and successful reactivation of the consolidated A memory at the start of Day 2. In group NA-A’BN-A, participants fully adapted to the opposing 30° CCW rotation (B) on Day 2, which was immediately washed out by no-perturbation trials (N) before it could consolidate. Despite B-learning being eliminated before consolidation, participants showed no savings for A on Day 3, relearning as naive on Day 3. In group NA-A’N-A, participants underwent the same reactivation followed immediately by N-trials, but without any intervening B-learning. In contrast to NA-A’BN-A, this group showed significant savings for A on Day 3, confirming that the A memory survived the washout intact. Insets: Average direction error (°) across the first three bins on Day 1 (red) and Day 3 (green) for each participant (dots), connected by lines to illustrate individual-level change. **(C)** Protection index for NA-A’BN-A and NA-A’N-A compared against group NA-A’B-NA from Experiment 1. The protection index of NA-A’BN-A was significantly lower than both NA-A’B-NA and NA-A’N-A, while NA-A’N-A did not differ from NA-A’B-NA. Individual participant values are shown as dots. **(D)** Direction error experienced at perturbation transitions on Day 1 (null-to-A) and Day 2 (B-to-null) in group NA-A’BN-A. The transition error at B-to-null is approximately 30° CW, the same direction as the original null-to-A transition on Day, leading the sensorimotor system to infer contextual similarity between the washout and the original A-learning episode, and consequently updating the A memory rather than B. **(E)** Direction error experienced at perturbation transitions on Day 1 (null-to-A) and Day 2 (A-to-null) in group NA-A’N-A. The transition error at A-to-null is approximately 30° CCW, opposite in direction to the original null-to-A transition, signaling a context change and leaving the A memory untouched. Together, panels D and E demonstrate that it is the directional relationship between successive prediction errors at perturbation transitions, rather than their magnitude alone, that determines which memory is updated.

To isolate the effect of N-trials on the reactivated A memory in the absence of any interfering rotation, we introduced a second group, *NA-A’N-A*. Participants successfully adapted to the 30° CW rotation (A) [*t*(9) = 30.99, *p* < 0.001, Cohen’s *d*_z_ = 10.33] on Day 1. Similar to *NA-A’BN-A*, participants successfully reactivated the consolidated A memory on Day 2 [*t*(9) = 1.19, *p* = 0.264, Cohen’s *d*_z_ = 0.40], evidence from Bayesian analysis supported this inference (*BF*_10_ = 0.55). This was immediately followed by N-trials, during which performance returned to baseline [*t*(9) = 14.52, *p* < 0.001, Cohen’s *d*_z_ = 4.84]. The critical question was whether this suppression reflected a temporary change in expression or an actual modification of the A memory trace, as indexed by savings on Day 3. Crucially, unlike *NA-A’BN-A*, participants in this group exhibited significant savings on Day 3 [*t*(9) = 7.77, *p* < 0.001, Cohen’s *d*_z_ = 2.59] (Figure 3B), indicating that A memory survived the Day 2 washout.

We then compared the protection indices of the two groups with that of the *NA-A’B-NA* group using a one-way ANOVA and found a significant group effect [*F*(2, 31) = 3.39, *p* = 0.046, ω² = 0.12]. Post-hoc comparisons confirmed no difference in the protection index of *NA-A’B-NA* and *NA-A’N-A* (*p* = 0.741), but there was a marginally significant difference in the protection index of *NA-A’BN-A* compared to *NA-A’B-NA* (*p* = 0.063) and *NA-A’N-A* (*p* = 0.084). This was further verified through Bayesian analysis, which provided supporting evidence for a difference in the protection index of *NA-A’BN-A* compared to *NA-A’B-NA* (*BF*_10_ = 1.95) and *NA-A’N-A* (*BF*_10_ = 2.78) (*Figure 3C*).

The contrasting outcomes between *NA-A’BN-A* and *NA-A’N-A* are particularly informative. In *NA-A’BN-A*, A memory was disrupted despite the interfering B rotation being washed out before consolidation, an outcome that cannot be accounted for by retrograde competition from a consolidated B memory. Instead, the N-trials themselves appear to have modified the A memory. We attribute this to the directional properties of the prediction error experienced at each perturbation transition. In *NA-A’BN-A*, the transition from B (30° CCW) to null generates an approximately 30° CW error, the same direction as the error experienced at the original null-to-A transition on Day 1 (Figure 3D). This directional match may have led the motor system to infer contextual similarity between the null session and the original A learning, resulting in inadvertent updating of the A memory rather than the B. In *NA-A’N-A*, by contrast, the transition from A (30° CW) to null generates an approximately 30° CCW error, which is directionally opposite to that experienced during the original null-to-A transition (Figure 3E). This mismatch signals a change in context, causing the N-trials to be encoded as a distinct experience that leaves the A memory untouched.

Together, the results of Experiments 1 and 2 provide convergent evidence that neither interference nor washout inherently disrupts a consolidated motor memory. Rather, the fate of the memory appears to be governed by the directional and contextual properties of the sensory prediction error experienced at each perturbation transition, which serve as a latent contextual cue driving memory protection or modification.

### Experiment 3: Sensory prediction error at perturbation transitions serves as a latent contextual cue for memory protection

The results from Experiments 1 and 2 suggest that sensory prediction error generated at the moment of perturbation transition might serve as a latent contextual signal that drives differentiation between the original and interfering memory representations. Experiment 3 was designed to test this hypothesis. If abrupt prediction error at the A’-to-B transition is indeed the critical signal for context differentiation, then eliminating that error by introducing the interfering rotation gradually rather than abruptly should abolish memory protection even when reactivation precedes interference. To test this, we introduced group *NA-A’Bgradual-NA*, in which the perturbation transitioned from A’ (30° CW) to B (30° CCW) across 120-trials in 0.5° increments, followed by 40-trials at the full 30° CCW rotation, thereby removing the large, abrupt prediction error that characterized the A’-to-B transition in Experiment 1.

Similar to Experiments 1 and 2, Day 1 began with a veridical-feedback baseline session followed by acquisition of the 30° CW rotation (A). Paired t-test revealed that Participants learned the A perturbation with a significant reduction of direction error from the early (Mean ± SD = 24.68 ± 2.49°) to the late (Mean ± SD = 7.08 ± 3.59°) phase of the learning session [*t*(10) = 15.57, *p* < 0.001, Cohen’s *d*_z_ = 4.92]. On Day 2, participants reactivated the consolidated A memory under 30° CW perturbed feedback, with end-of-reactivation direction error not differing significantly from the last bin of Day 1 learning [*t*(10) = 0.89, *p* = 0.391, Cohen’s *d*_z_ = 0.28], confirming successful reactivation with strong support for the null hypothesis from Bayesian analysis (*BF*_10_ = 0.42). The gradual introduction of the interfering B rotation then produced partial adaptation, with direction error changing significantly from early (Mean ± SD = 3.87 ± 4.21°) to late (Mean ± SD = −15.32 ± 6.47°) phase of the interference session [*t*(10) = 7.32, *p* < 0.001, Cohen’s *d*_z_ = 2.31].

On Day 3, following a washout session to eliminate anterograde effects of B-learning, participants relearned the A rotation. In contrast to Experiment 1, no savings were observed [*t*(10) = 2.16, *p* = 0.056, Cohen’s *d*_z_ = 0.68] (Figure 4A). We next compared its protection index with *NA-A’B-NA*. Because Levene’s test indicated a significant violation of the equal-variance assumption [*F*(1, 21) = 6.51, *p* = 0.02], Welch’s t-test was used in place of the standard Student’s t-test. The protection index of *NA-A’Bgradual-NA* was significantly lower than that of *NA-A’B-NA* [Welch’s *t*(18.27) = 2.39, *p* = 0.028, Cohen’s *d* = 1.12] (Figure 4B). Despite reactivation having preceded the interfering rotation, the same sequence that protected A memory in Experiment 1, the absence of an abrupt prediction error at the transition rendered reactivation insufficient to protect the original memory. This was confirmed by comparing the change in the error experienced when participants transitioned from A’ to B (61.42°) and from A’ to B (gradual) perturbation (0.67°) (Figure 4C). In the case of A’ to Bgradual perturbation the experienced error was similar to zero (0.67°). When the motor system received no signal to infer a context change, the gradually introduced B rotation updated the existing A memory rather than being encoded as a separate representation.

**Figure 4.**
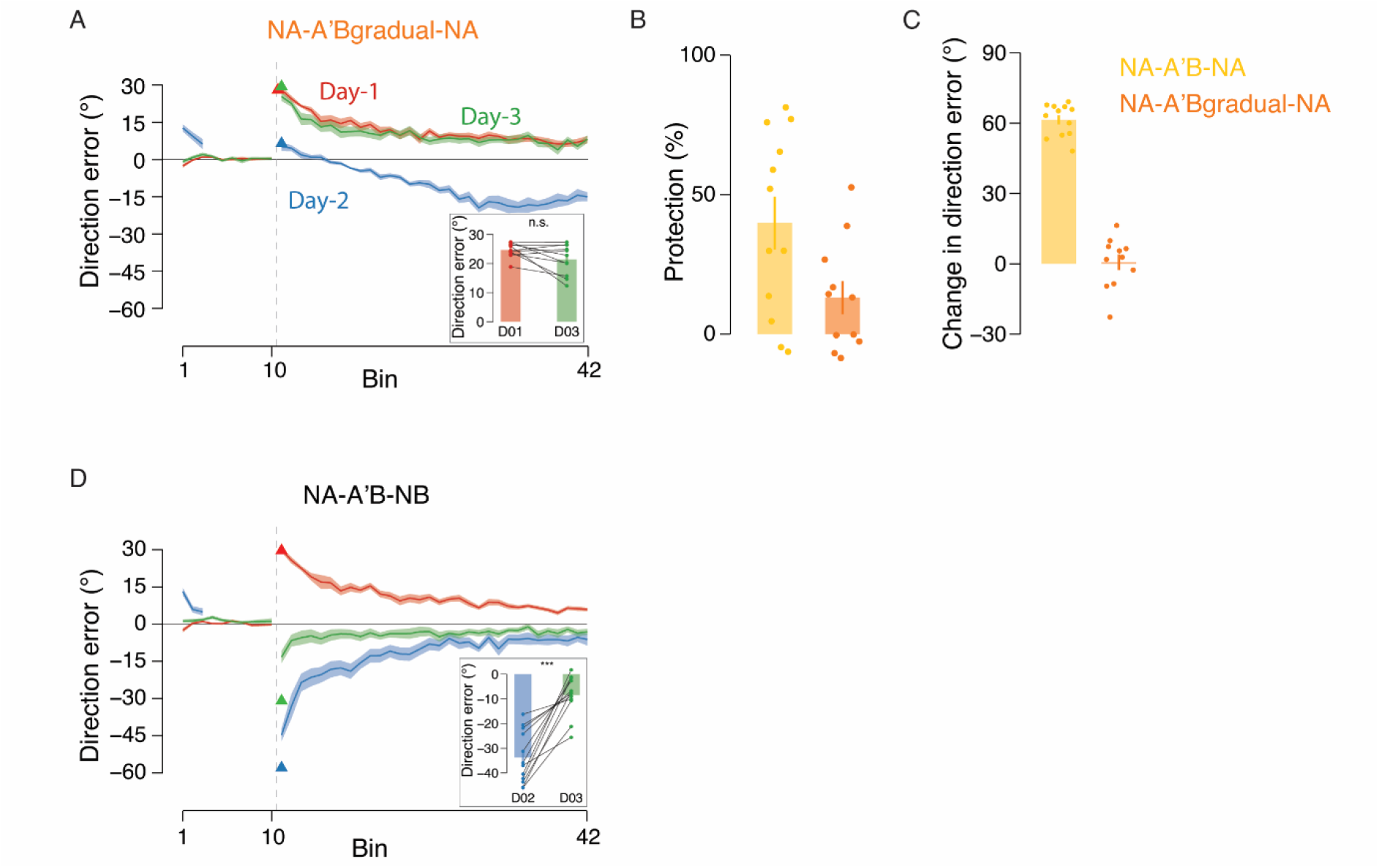
An abrupt sensory prediction error at the perturbation transition is necessary for memory protection, and its presence supports the independent consolidation of the interfering rotation as a separately retrievable memory representation. Change in mean direction error (°) across bins for participants in **(A)** group NA-A’Bgradual-NA and **(D)** group NA-A’B-NB. Shaded ribbons indicate ± SE across participants. Triangles mark the first trial of the Day 1 learning, Day 2 interference, and Day 3 relearning sessions. Red, blue, and green colors denote Day 1, 2, and 3 respectively. Both groups showed successful acquisition of the 30° CW rotation (A) on Day 1 and successful reactivation of the consolidated A memory at the start of Day 2. In group NA-A’Bgradual-NA, the interfering 30° CCW rotation (B) was introduced gradually in 0.5° increments across 120-trials and was then held constant for 40-trials at 30° CCW rotation, eliminating the abrupt prediction error that characterized the A’-to-B transition in Experiment 1. Despite identical reactivation and adaptation to the interfering task B, participants showed no savings for A on Day 3, relearning it at a rate indistinguishable from that of naive Day 1 acquisition. This demonstrates that reactivation alone is insufficient to protect the consolidated memory when the transition prediction error is removed; it is the abrupt context-change signal, not the retrieval event itself, that drives separate encoding of the interfering rotation. In group NA-A’B-NB, participants underwent the identical Day 2 procedure as NA-A’B-NA from Experiment 1, but were retested on B rather than A on Day 3. Participants showed significant savings for B despite an intervening washout, confirming that B was independently consolidated as a distinct memory rather than simply failing to overwrite A. Insets: Average direction error (°) across the first three bins on Day 1 (red) and Day 3 (green) for each participant (dots), connected by lines to illustrate individual-level change. **(B)** Protection index for NA-A’Bgradual-NA compared against NA-A’B-NA from Experiment 1. The protection index of NA-A’Bgradual-NA was significantly lower than that of NA-A’B-NA, confirming that the gradual elimination of the transition prediction error abolished the memory protection conferred by reactivation. Individual participant values are shown as dots. **(C)** Direction error experienced at the A’-to-B transition on Day 2 for groups NA-A’B-NA (abrupt) and NA-A’Bgradual-NA (gradual). The abrupt transition generated a large single-trial prediction error (∼61°), whereas the gradual transition produced a near-zero change in error per trial (∼0.67°), confirming that the gradual schedule effectively eliminated the context-change signal available to the sensorimotor system. Together, panels A–C provide evidence that abrupt prediction error at the perturbation transition is necessary for context differentiation, while panel D shows that, when that signal is present, the interfering rotation can be consolidated independently and remains retrievable as a distinct memory representation.

These findings provide supporting evidence that the memory protection observed in Experiments 1 and 2 is not a consequence of reactivation per se but depends specifically on the context change driven by the abrupt sensory prediction error generated at the perturbation transition. A critical corollary of this conclusion is that when such a prediction error is present, as in Experiment 1, the interfering B rotation could be encoded as a genuinely distinct memory rather than merely failing to overwrite A. If this is correct, B memory should be independently retrievable following the Day 2 session.

To test this possibility, we included the *NA-A’B-NB* control group, which underwent the identical Day 2 procedure as *NA-A’B-NA* (reactivation of A followed by full B-learning) but was retested on B rather than A on Day 3 following washout. Participants demonstrated significant savings for B [*t*(11) = 6.84, *p* < 0.001, Cohen’s *d*_z_ = 2.06] (Figure 4D), confirming that B was retained as an independent memory despite the intervening washout. Crucially, this finding demonstrates that B was independently consolidated and remained retrievable following the Day 2 session, consistent with context-dependent memory segmentation. (Bouton, 1993; Heald et al., 2021).

Taken together, Experiments 1-3 and the *NA-A’B-NB* control group support an important role for abrupt transition-related prediction error as a latent contextual signal for motor memory protection, and suggest that its consequence is not merely interference resistance but the independent consolidation of competing rotations as separately retrievable memory representations. In Experiment 4, we explore whether an explicit contextual cue, independent of any sensory prediction error, can achieve the same protective effect.

### Experiment 4: Explicit contextual cues protect consolidated motor memories independent of sensory prediction error

We have established that memory protection following interference is contingent on the availability of a signal that informs the motor system of a contextual change. An error-based cue generated at the moment of perturbation transition allows the motor system to infer that the interfering experience arises from a distinct context. In the previous experiments, this signal is latent and emerges implicitly from the sensorimotor consequences of transitioning between perturbations. In Experiment 4, we asked whether an explicit contextual cue uniquely associated with each perturbation could confer the same protective effect, and, critically, whether such a cue would protect memory even in conditions where there is no transition (B-learning without reactivation on Day 2), thereby leading to retrograde interference in Experiment 1. For this, we used the groups *NA-B-NA* and *NA-A’B-NA*, similar to experiment 1, but we tagged the two perturbations with distinct cues as a secondary follow-through target.

As in the previous 3 Experiments, both the groups completed a veridical-feedback baseline session on Day 1 followed by acquisition of the 30° CW (A) perturbation. A mixed ANOVA showed a significant reduction in the direction error from early to late sessions [*F*(1, 18) = 67.30, *p* < 0.001, ω² = 0.58] and equivalent acquisition of A with no significant between-group difference [*F*(1, 18) = 1.63, *p* = 0.218, ω² < 0.01] confirmed with support against group effect from Bayesian analysis (*BF*_10_ = 0.62).

On Day 2, participants in the *Context-NA-A’B-NA* group reactivated the consolidated A memory before encountering the interfering B rotation. Participants reactivated the memory in the presence of 30° CW (A) rotation over 15-trials, and there was no significant difference in the last bin of learning and reactivation [*t*(9) = 0.40, *p* = 0.698, Cohen’s *d*_z_ = 0.13], confirmed with strong support for the null hypothesis from Bayesian analysis (*BF*_10_ = 0.33). Both *Context-NA-A’B-NA* (following reactivation) and *Context-NA-B-NA* then fully adapted to the opposing 30° CCW rotation across the interference session [*F*(1, 18) = 151.85, *p* < 0.001, ω² = 0.64], reaching equivalent asymptotic performance with no significant between-group difference at the end of B training [*F*(1, 18) = 1.51, *p* = 0.236, ω² < 0.01] with support against group effect from Bayesian analysis (*BF*_10_ = 0.71).

On Day 3, both groups completed a washout session to eliminate the anterograde effects of B-learning before relearning A. In contrast to the no-context *NA-B-NA* group from Experiment 1, which showed no savings, both *Context-NA-B-NA* [*t*(9) = 7.42, *p* < 0.001, Cohen’s *d*_z_ = 2.47] (Figure 5A) and *Context-NA-A’B-NA* [*t*(9) = 3.49, *p* = 0.007, Cohen’s *d*_z_ = 1.16] (Figure 5B) showed significant savings during A relearning. A one-way ANOVA comparing the protection index of the two context groups against the no-context *NA-A’B-NA* group from Experiment 1 revealed no significant difference across the three groups [*F*(2, 29) = 0.21, *p* = 0.81, ω² = 0.05] (Figure 5E) with support for the null hypothesis from Bayesian analysis (*BF*_10_ = 0.24). The equivalent degree of memory protection achieved by explicit contextual cueing and implicit prediction-error-based cueing indicates that contextual inference, rather than the specific form of the contextual signal, is the governing mechanism underlying memory protection.

**Figure 5.**
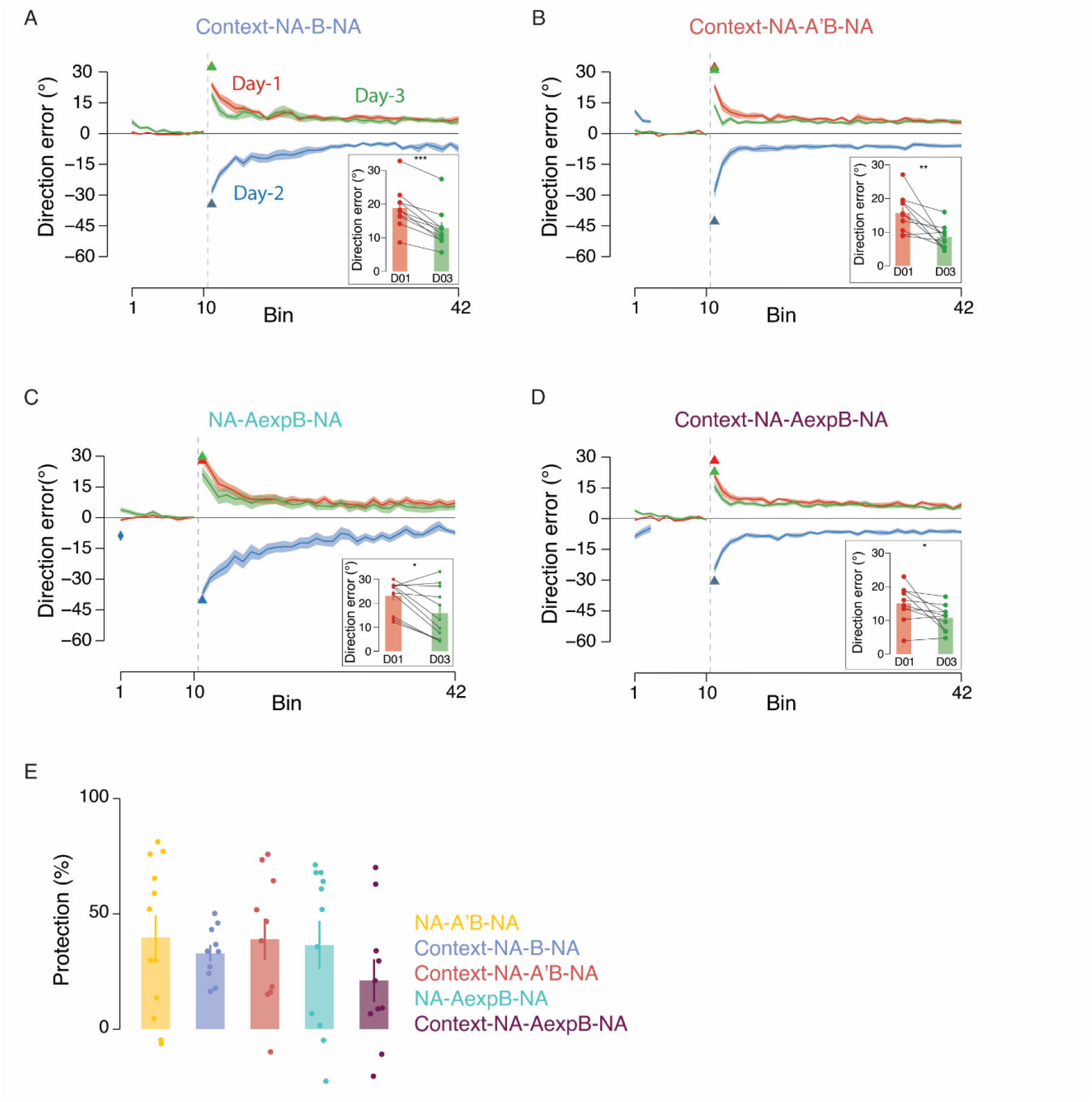
Explicit contextual cues and error-free expression of the consolidated motor memory each independently protected it against interference. Change in mean direction error (°) across bins for participants in **(A)** group Context-NA-B-NA, **(B)** group Context-NA-A’B-NA, **(C)** group NA-AexpB-NA, and **(D)** group Context-NA-AexpB-NA. Shaded ribbons indicate ±SE across participants. Triangles mark the first trial of the Day 1 learning, Day 2 interference, and Day 3 relearning sessions. Red, blue, and green colors denote Day 1, 2, and 3 respectively. All four groups successfully acquired the 30° CW rotation (A) on Day 1 and fully adapted to the opposing 30° CCW rotation (B) on Day 2, achieving equivalent asymptotic performance across groups within each experiment. In Experiment 4 (panels A and B), the two rotations were tagged with distinct explicit contextual cues in the form of a secondary follow-through target. Group Context-NA-B-NA received B-learning on Day 2 without any prior reactivation of A, the condition that produced complete retrograde interference in the no-cue NA-B-NA group of Experiment 1. Despite the absence of reactivation, participants showed significant savings for A on Day 3, demonstrating that the explicit cue alone was sufficient to drive separate encoding of B and protect the original A memory without requiring any prior memory access. Group Context-NA-A’B-NA reactivated A before B-learning, as in Experiment 1, and similarly showed robust savings on Day 3. In Experiment 5 (panels C and D), the A memory was expressed prior to B-learning in the complete absence of error feedback. Both groups showed significant savings for A on Day 3, with protection indices statistically equivalent to those of the NA-A’B-NA group from Experiment 1. This demonstrates that error-driven retrieval is not a prerequisite for memory protection: memory expression is sufficient to re-establish the A-context motor state from which the abrupt transition prediction error is generated at the onset of B-learning, and that transition error alone is sufficient to drive context differentiation. Insets: Average direction error (°) across the first three bins on Day 1 (red) and Day 3 (green) for each participant (dots), connected by lines to illustrate individual-level change. **(E)** Protection index compared across all four groups and the no-cue NA-A’B-NA group from Experiment 1. No significant differences in protection index were observed across any of the groups, establishing that explicit contextual cueing, error-free expression, and error-driven reactivation all confer equivalent memory protection, and that the governing variable across all conditions is the availability of a reliable context-change signal at the moment interference is introduced rather than the specific mechanism through which that signal is generated. Individual participant values are shown as dots.

The results of Experiment 4 extend the contextual inference framework established across Experiments 1–3 in two important respects. First, the identity of the contextual signal does not determine whether protection occurs: an explicit environmental cue and an implicit sensory prediction error result in equivalent protection, indicating that what matters is the availability of any reliable signal that allows the motor system to tag the interfering experience as arising from a distinct context. Second, and more consequentially, memory protection does not require prior reactivation of the original memory. In Experiment 1, the absence of reactivation before B-learning (*NA-B-NA*) produced complete retrograde interference; here, the presence of an explicit cue alone was sufficient to prevent that interference, even without any prior access to the A memory. This dissociation demonstrates that the protective function previously attributed to reactivation in Experiment 1 was not intrinsic to retrieval itself, but arose because reactivation generated the prediction error that served as an implicit context-change signal, a function that an explicit cue can perform independently. Together, Experiments 1–4 converge on a single organizing principle: motor memory protection and disruption following interference are governed by whether the motor system can infer a change in context at the moment the interfering perturbation is introduced. When such a signal is available, whether latent or explicit, the interfering experience is encoded as a new memory representation, leaving the original intact. When it is absent, the interfering perturbation updates and overwrites the existing memory.

### Experiment 5: Is error-driven retrieval necessary for memory reactivation?

So far, we have shown that memory protection or modification is guided by contextual inference, and that the contextual signal can be either an implicit sensory prediction error or an explicit environmental cue, Experiment 5 addresses a more targeted mechanistic question: is error-driven retrieval of the consolidated memory necessary to generate a sensory prediction error at the perturbation transition, or is expression of that memory in the complete absence of error feedback equally sufficient?

To test this, we introduced the group *NA-AexpB-NA*, in which participants expressed the consolidated A memory on Day 2 through five reaching movements performed without cursor feedback, thereby eliminating error-driven retrieval while preserving motor expression. As in previous experiments, Day 1 began with a veridical-feedback baseline session followed by acquisition of the 30° CW rotation (A), confirmed by significant error reduction from early to late learning [*t*(10) = 8.52, *p* < 0.001, Cohen’s *d*_z_ = 2.69]. On Day 2, participants successfully expressed the A memory during the no-feedback session, with hand angles significantly different from zero [*t*(10) = 3.94, *p* < 0.01, Cohen’s *d* = 1.25], following which they successfully adapted to the interfering B perturbation [ t(10) = 9.94, *p* < 0.001, Cohen’s *d*_z_ = 3.14]. Similar to NA-AB-NA, participants exhibited significant savings for A learning on Day 3 [*t*(10) = 3.15, *p* = 0.01, Cohen’s *d*_z_ = 1.00] (Figure 5C).

To examine whether the same result holds when expression is combined with an explicit contextual cue, we also introduced the group *Context NA-AexpB-NA*, in which participants were exposed to 15 error-clamped A perturbation trials on Day 2, tagged with an explicit cue (secondary follow-through target) to distinguish the perturbation. Participants adapted to 30° CW rotation A [*t*(9) = 4.29, *p* = 0.002, Cohen’s *d*_z_ = 1.43] on Day 1, and showed a significant reduction in direction error from first to last expression bin [*t*(9) = 2.83, *p* = 0.019, Cohen’s *d*_z_ = 0.94]. Following the expression session, participants successfully adapted to the interfering perturbation [*t*(9) = 7.27, *p* < 0.001, Cohen’s *d*_z_ = 2.42]. Participants showed significant savings during the relearning session on Day 3 [*t*(9) = 2.45, *p* = 0.036, Cohen’s *d*_z_ = 0.82] (Figure 5D). A one-way ANOVA comparing the protection index of NA-AexpB-NA and *Context-NA-AexpB-NA* against *NA-A’B-NA* from Experiment 1 revealed no significant difference across the three groups [*F*(2, 30) = 0.99, *p* = 0.38, ω² < 0.01] (Figure 5E) with Bayesian analysis confirming support for the null hypothesis (*BF*_10_ = 0.38). The equivalent protection across groups that differed in whether expression was error-driven or error-free indicates that error-based retrieval is not a prerequisite for memory protection, and that expression of memory in the absence of error feedback is sufficient to confer memory protection, consistent with the interpretation that error-free expression re-establishes the A-context motor state from which a transition prediction error is generated at the onset of B, though the transition error itself was not directly measured in these groups.

The mechanism underlying this equivalence lies in the structure of the perturbation transition itself. Regardless of whether the A memory is accessed through error-driven reactivation or error-free expression, the abrupt shift from A context motor output to the opposing B context generates a large sensory prediction error at the transition point, the same signal that drove context differentiation in Experiments 1 and 3. It is this transition-point prediction error, not the form of prior memory access, that informs the motor system of a context change and triggers encoding of B as a separate memory trace. Taken together, the results of Experiment 5 establish that the boundary condition for memory protection is not the retrieval mechanism employed, but the availability of a sufficiently large prediction error at the moment interference is introduced.

## DISCUSSION

How does the brain decide whether to update an existing memory or create a new one? This question sits at the intersection of memory stability and plasticity, which is the fundamental tension in any biological system that must both preserve acquired knowledge and remain sensitive to change. In the domain of motor memory, the question takes a specific form: when an opposing perturbation follows the consolidation of a learned adaptation, does it overwrite the original, and if not, what prevents it? In the present study, we investigated how interfering information can affect already consolidated motor memories and what guides this process. Although consolidated memories were once thought to be resistant to any interference (McGaugh, 2000; Squire & Alvarez, 1995), accumulating evidence over the past two decades has shown that memory reactivation returns them to a transiently labile state during which they are susceptible to modification or disruption before reconsolidation (Lee et al., 2017; Nader et al., 2000; Walker et al., 2003). Across five experiments, we show that the fate of a consolidated motor memory under interference is not determined by reactivation, interference, or washout per se, but is predicted by the availability of a contextual signal at the moment interference is introduced. When such a signal was available, whether as an implicit sensory prediction error or as an explicit environmental cue, the interfering experience was encoded as an independent memory trace, leaving the original intact. When no such signal was available, the interfering experience produced greater disruption of the original memory, reflected in the absence of savings. These findings are consistent with a contextual inference account and may help resolve a long-standing inconsistency in the motor reconsolidation literature.

### Resolving an inconsistency in the reconsolidation literature

The discovery that reactivation renders consolidated memories transiently labile (Nader et al., 2000) offered an alternative framework to the classical view that consolidated engrams are permanent and fixed (McGaugh, 2000; Squire & Alvarez, 1995). This was confirmed in human motor memory where training on a second motor sequence immediately after reactivating the first produced retrograde interference, whereas the same training without prior reactivation did not (Walker et al., 2003). Subsequent work, however, did not converge on this evidence. Several studies reported that interference introduced after reactivation could strengthen rather than impair (Herszage & Censor, 2017; Wymbs et al., 2016) or fail to disrupt (Hardwicke et al., 2016) the original memory, challenging the prediction that reactivation makes the memory labile and vulnerable. Furthermore, retrograde interference has also been reported in the complete absence of reactivation (Caithness et al., 2004; Krakauer et al., 2005). No single principle has explained why reactivation sometimes leads to protection, other times to disruption, and in some cases not even necessary for disruption to occur.

Our data might resolve this inconsistency by showing that the operative variable is not reactivation but the contextual signal generated at the transition into the interfering learning session. Reactivation followed by an abrupt interfering rotation protected the original memory; the identical rotation introduced without reactivation did not. Critically, the responsible mechanism is not reactivation-based destabilization (Nader et al., 2000), rather it is the abrupt prediction error at the transition that determines whether the interfering experience is distinguished from the existing memory, resulting in the formation of a new trace or is inferred as the existing one, leading to its modification. Reactivation matters only because it establishes the baseline motor state from which that transition error can be measured. The disruption observed without reactivation (Caithness et al., 2004; Krakauer et al., 2005) could be explained as competitive interference between two adaptations sharing an undifferentiated context. Protection and disruption are therefore not opposing outcomes of retrieval but rather two outcomes of a single computation, contextual inference, tipped in either direction by whether a context-change signal is present. Prior work in the visuomotor adaptation domain has generated findings that appear mutually contradictory but are coherently explained by the contextual inference framework. Studies which introduced an opposing rotation directly without prior reactivation and found retrograde interference observed as participants relearning the original rotation as naive (Caithness et al., 2004; Krakauer et al., 2005). Under the present account, the absence of any prior memory access meant no transition prediction error was available to signal a context change; the two rotations might compete within a shared representational context, and the interfering rotation dominated. Conversely, when participants were exposed to subtle sensorimotor variability around the original task immediately after reactivation, such that the intervention remained within the original learning context, strengthening rather than disruption was observed. One interpretation consistent with the present framework is that modifications were small enough that no reliable context-change signal was generated at the reactivation-to-intervention boundary. As a result, the motor system updated the existing trace (strengthening it) rather than encoding a new one. However, the transition prediction errors were not measured in this study (Wymbs et al., 2016). Finally, the failure to find reliable disruption following post-reactivation interference in a sequence-learning paradigm (Hardwicke et al., 2016) remains unexplained under the reactivation-vulnerability account. One speculative interpretation, consistent with the present framework but not tested by the authors is that prediction error between two distinct sequences accumulates gradually across trials as the new sequence is discovered, rather than manifesting as a concentrated single-trial transition signal as in visuomotor rotation, leaving the motor system without an unambiguous context-change cue, and producing neither reliable protection nor reliable disruption. These three contrasting outcomes, disruption without a contextual signal, strengthening through within-context updating, and null results with distributed error, form a coherent pattern under the contextual inference account.

### Prediction error as a causal, direction-sensitive contextual signal

The conceptual link between prediction error and memory updating has a long history in both computational modelling and memory reconsolidation research. Rescorla and Wagner’s foundational learning rule formalized the idea that learning is driven by the discrepancy between expected and obtained outcomes (Rescorla & Wagner, 1972). Evidence from classical conditioning paradigm supports this. Reactivating the fear memory in humans triggered reconsolidation and left the trace vulnerable to pharmacological disruption only when the reactivation episode contained a prediction error mismatch; when reactivation was predictable and error-free, the memory was expressed but not destabilized (Sevenster et al., 2013). This is further supported by evidence from episodic memory domain, which show that the surprise elicited by an incomplete reminder determined whether the reactivated memory was susceptible to reconsolidation-based distortion (Sinclair & Barense, 2018, 2019). Collectively, these studies established a boundary condition: prediction error at retrieval governs whether reactivation opens a reconsolidation window.

The present study demonstrates a functionally distinct but mechanistically related role for prediction error, not at retrieval, but at the moment interference is introduced. These two roles of prediction error, as a driver of context differentiation in the COIN model, and as a trigger for reconsolidation-mediated updating, operate at different points in the memory episode and are not necessarily in conflict. The Experiment 3 manipulation is the critical piece of evidence. By introducing the interfering rotation in 0.5° increments across 120-trials, we eliminated the abrupt transition error while preserving identical reactivation, identical session structure, and learning of the interfering rotation. Participants did not show any savings. This could be due to the absence of the transition error during the transition from reactivation to the interfering learning. We note, however, that the gradual introduction produced only partial rather than complete adaptation, and this partial-adaptation confound cannot be fully ruled out in the present design.

More surprisingly, and more theoretically consequential, is the finding from Experiment 2 that the direction of the prediction error, not solely its magnitude, determines which memory is updated. When an immediate washout session following B-learning generated a transition error in the same rotational direction as the original null-to-A acquisition error (approximately 30° clockwise), the motor system updated the A memory rather than the B, producing disruption despite B undergoing washout before it could consolidate. When the same washout procedure was applied following A reactivation alone, the A-to-null transition generated an error in the opposite direction (approximately 30° counterclockwise), which the motor system correctly interpreted as a context change and left A intact. The magnitude of the prediction error was comparable in both cases; only its direction differed. This directional specificity might imply that contextual inference in the motor system is not simply a threshold on the magnitude of prediction error; it could compute the relationship between successive prediction errors across transitions, incorporating their directional structure. We offer this as a hypothesis generated by the data rather than a conclusion it directly supports. A prospective test would require parametrically varying the direction and magnitude of the transition error independently, while holding all other features of the learning constant, to establish whether direction is indeed the operative dimension. If confirmed, this would constitute a meaningful refinement of scalar surprise-based accounts of context inference, including the current COIN model formulation (Heald et al., 2021). Whether the motor system encodes a signed or vector-valued prediction error at perturbation transitions, and how that signal maps onto the known directional tuning of cerebellar climbing fiber inputs (Shadmehr et al., 2010), is a question for future neurophysiological and computational investigation.

### Context differentiation produces independently retrievable traces

A further prediction of the contextual inference account is that successful context differentiation should support the independent consolidation of the interfering experience as a distinct trace, rather than merely sparing the original from disruption. We tested this directly by retesting the memory of the interfering rotation itself in participants who had reactivated the original memory beforehand. The interfering rotation was retained as savings despite an intervening washout, demonstrating that it had been consolidated as an independent representation rather than merely failing to overwrite the original. This result rules out the parsimonious possibility that the transition error simply prevented the interfering rotation from being learned at all.

This finding, along with savings for A observed in the *NA-A’B-NA* group, indicates that when reactivation precedes abrupt B-learning, both the original and the interfering memory can independently survive to Day 3. Because A and B retention were tested in separate groups of participants, we cannot directly conclude that the two memories coexisted simultaneously within the same individual. The data establish that neither memory was eliminated by the Day 2 session and that each remained independently accessible following an intervening washout, a pattern consistent with context-dependent memory segmentation, though the stronger claim of parallel consolidation would require a within-subject test.

This outcome resonates with Bouton’s (1993) account of extinction and renewal, in which extinction does not erase the original conditioned response but creates a competing memory whose expression is context-gated (Bouton, 1993). The present results suggest an analogous structure may operate in motor memory: rather than one rotation overwriting the other, context differentiation may support the formation of two separable representations whose independent retrievability is preserved.

### Explicit cues and error-free expression converge on the same mechanism

Two further manipulations clarify what form the contextual signal must take, and how the original memory must be accessed for it to be effective. First, tagging the original and interfering rotations with distinct explicit cues protected the original memory even without any reactivation preceding interference. This demonstrates that reactivation’s protective role in the uncued condition was purely instrumental: it mattered only because it generated a prediction error at the subsequent transition, a function that an explicit cue can perform without any retrieval at all. This situates the present work within a broader family of occasion-setting accounts of context-gated memory expression, including extinction and renewal (Bouton & Bolles, 1979; Bouton & Ricker, 1994) and source-context effects in episodic memory, where a subtle reminder of the original context determines whether new information is integrated or kept separate (Hupbach et al., 2007). Contextual information, whether sensory, associative, or environmental, appears to function as a general-purpose gating signal for memory updating across these domains, rather than as a mechanism unique to motor memory.

Second, expressing the consolidated original memory without any corrective feedback protected it from interference just as effectively as error-driven reactivation. This could be because the critical prediction error arises at the transition into the interfering rotation rather than during the preceding retrieval episode. This is consistent with forward-model accounts of motor control, in which a predicted sensory consequence is generated for any planned movement, regardless of whether corrective feedback follows (Wolpert & Kawato, 1998), and indicates that reactivation-based destabilization of the original memory is not a necessary precondition for context differentiation at the subsequent transition.

### Limitations

The present findings should be interpreted in light of several limitations. The present paradigm does not neurally dissociate reconsolidation-mediated updating from simple competitive interference in the uncued, non-reactivated condition; distinguishing them will require markers specific to the reconsolidation process itself. The savings measure used throughout may conflate genuine implicit adaptation with explicit strategy reuse; strategy-exclusion designs would yield a cleaner index of implicit retention. Relatedly, explicit aiming strategies were not measured across experiments. Given that a 30° rotation is large enough to invite deliberate re-aiming, a portion of the savings observed on Day 3 may reflect retrieval of a previously effective explicit strategy rather than retention of the implicit motor adaptation; future designs incorporating process-dissociation procedures or reaction-time probes would clarify the relative contribution of each. The directional account of the washout finding was developed after the result was observed, and a parametric test that independently manipulates transition-error sign and magnitude is needed to confirm it. Finally, group sizes were modest, and the sample was restricted to healthy young adults, leaving open the question of whether these mechanisms generalize across the lifespan or to populations with hippocampal or cerebellar impairment.

### Conclusion

This work demonstrates that the fate of consolidated motor memory under interference is determined by contextual inference rather than by reactivation itself. Sensory prediction error at the moment of contextual transition or explicit environmental cues was sufficient to support the independent consolidation of competing motor memories as separately retrievable representations. The directional structure of prediction error about how the sign of the transition error relative to prior learning history may determine which memory is targeted is an additional hypothesis generated by these findings that requires future confirmation. By identifying the mechanistic signal that governs protection versus disruption, this study unifies a previously inconsistent literature on motor memory reconsolidation and extends a general computational principle of context-dependent memory formation, established across fear conditioning and episodic memory, into the domain of human motor learning.

## ACKNOWLEDGEMENT

We thank IIT Hyderabad for all the institutional-level support.

## DATA AVAILABILITY

The datasets used and/or analyzed during the current study are available from the corresponding author upon reasonable request.

